# Early-life paternal relationships predict adult female survival in wild baboons

**DOI:** 10.1101/2025.01.22.634342

**Authors:** David A.W.A.M. Jansen, J. Kinyua Warutere, Jenny Tung, Susan C. Alberts, Elizabeth A. Archie

**Affiliations:** Department of Biological Sciences, University of Notre Dame, Notre Dame, IN, USA; Department of Pathobiological Science, School of Veterinary Medicine, University of Wisconsin-Madison, United States; Amboseli Baboon Research Project, Amboseli National Park, Kajiado, Kenya; Department of Evolutionary Anthropology, Duke University, Durham, NC, USA; Department of Biology, Duke University, Durham, NC, USA; Duke University Population Research Institute, Duke University, Durham, NC, USA; Department of Primate Behavior and Evolution, Max Planck Institute for Evolutionary Anthropology, 04103 Leipzig, Germany; Canadian Institute for Advanced Research, Toronto, Canada M5G 1M1, Canada; Faculty of Life Sciences, Institute of Biology, Leipzig University, Leipzig, Germany

**Keywords:** paternal care, parental care, mammals, early-life effects, fathers, adult survival

## Abstract

Parent-offspring relationships can have profound effects on offspring behavior, health, and fitness in adulthood. These effects are strong when parents make heavy investments in offspring care. However, in some mammals, including several species of carnivores, rodents, and primates, fathers live and socialize with offspring, but paternal care *per se* is subtle or indirect. Do these limited father-offspring relationships also affect later-life outcomes for offspring? Working in a well-studied baboon population where males contribute little direct offspring care, we found that juvenile female baboons who had stronger paternal relationships, or who resided longer with their fathers, led adult lives that were 2-4 years longer than females with weak or short paternal relationships. This pattern did not differ between females who experienced high versus low levels of early-life adversity; hence, paternal relationships were equally protective in both harsh and benign early environments. Males’ relationships were strongest with juvenile females they were most likely to have sired and when males had few mating opportunities. Hence, father-daughter relationships may be constrained by male mating effort. Because survival predicts female fitness, fathers and their daughters may experience selection to engage socially and stay close in daughters’ early lives.

## 1. INTRODUCTION

In humans and other mammals, social environments are powerful determinants of individual health, survival, and fitness [1]. Social relationships in early life are especially important, both because of their immediate benefits to offspring—such as opportunities to learn social skills, gain resources, or receive protection—and also because these relationships have lasting consequences for adult health and survival [2]. Maternal relationships are especially well-studied in this regard [3–11], and across mammals, maternal loss and the quality of maternal care have lasting consequences for offspring gene regulation, stress reactivity, social integration, and adult survival [3–12].

But what about relationships with fathers? Early-life paternal social effects have received less attention, in part because it is rare for male mammals to make substantial investments in offspring care [13, 14]. However, in species with caring males, early-life father-offspring relationships can have profound effects on offspring in adulthood. In humans, paternal absence in childhood is linked to lower income, poorer health, and higher mortality risk [e.g., 15, 16–19]. In rodents with biparental care, fathers affect the complexity of offspring social environments, with consequences for neurological development and adult behavior [20, 21]. Yet mammal species with caring males are rare. In a wider (but still unusual) set of group-living mammals, fathers live and even socialize with offspring, but paternal care is subtle and often indirect [13, 22]. These species include several carnivores and equids, as well as gorillas, chimpanzees, baboons, and other primates [13, 17, 23–29]. Whether these more limited early-life paternal relationships have long-term consequences for offspring is largely unknown.

Here we test if early-life paternal relationships predict adult survival for female baboons—a species where fathers and their juvenile offspring may co-reside and interact, but where mothers provide all essential care [30, 31]. Baboons are useful for testing these relationships for three reasons. First, many baboons live in polygynandrous mating systems where paternity certainty is incomplete, yet adult males often interact with their offspring and engage in some forms of offspring care, including carrying and supporting offspring in conflicts [30, 32–38]. Further, lactating female baboons sometimes form close social bonds (i.e., “primary associations”) with particular males, and these relationships are better explained by parenting than mating effort [33–35, 39–41], as male primary associates typically do not sire their female associate’s next infant [34, but see 35]. Male primary associates are also disproportionately the fathers of their female partner’s current infant, intervene on behalf of females and their infants in conflicts, and may buffer infants from rough handling [30, 32–38, 42].

Second, proximity to adult males and/or paternal presence in early life have developmental and social consequences for young baboons. For instance, proximity to adult males, including fathers and non-fathers, increases the complexity of the social environment for infants [43]. Paternal presence is also correlated with earlier sexual maturity in daughters [32] and predicts stronger social bonds between paternal half-siblings [44].

Third, adult lifespan, our outcome of interest, explains 80%-90% of the variance in lifetime reproductive success for female baboons in our population [7, 45, 46]. Hence, if early-life paternal relationships influence daughters’ lifespans, they are likely to have important consequences for fitness. Males and their daughters may therefore experience selection to form and maintain early-life social relationships with one another, and male care may meet the criteria for “true” parental care (i.e., care that improves offspring and male fitness [14]).

Working in the Amboseli baboon population in Kenya [47], we pursued three objectives to understand whether and how fathers influence daughters’ lifespans. First, we measured patterns of grooming and co-residency between juvenile females and their fathers. Social bonds are often developed and maintained through grooming, a primary affiliative behavior in many social species, including baboons [48–50]. We measured co-residency because father-offspring pairs who live together for longer have more time to interact (co-residency varies because males may disperse or die during their daughters’ juvenile years).

Second, we tested whether juvenile females who had stronger grooming relationships or longer co-residency with their fathers exhibit higher adult survival than females who had weak paternal grooming relationships or short co-residency periods. In the Amboseli baboons, adult female longevity is also predicted by an accumulation of harsh conditions in early life, including drought, maternal loss, or having a low-ranking or socially isolated mother [7, 45, 51]. Hence, we also tested if early-life relationships with fathers protect daughters from the negative effects of cumulative early-life adversity.

Third, we tested why some fathers are more likely to groom or have longer co-residency with their daughters than others. We predicted that males would have stronger relationships with their daughters when they had high paternity certainty (e.g., spent more time mate guarding his daughter’s mother when his daughter was conceived) and when reproductive tradeoffs were favorable (e.g., when the male had few current mating opportunities). Together, our results support the importance of early-life paternal relationships to adult female baboons, lending context to paternal effects on adult outcomes and the evolution of mammalian parental care.

## 2. METHODS

### (a) Study population and subjects

Our subjects were wild baboons studied by the Amboseli Baboon Research Project (ABRP) in the Amboseli ecosystem, Kenya [47]. This population is admixed between yellow and anubis baboons (*Papio cynocephalus* and *P. anubis*), with majority yellow ancestry [52, 53]. ABRP observers collect behavioral and demographic data year-round on a near-daily basis, and all study animals are known through visual recognition. Our analyses centered on 216 female baboons that: (i) survived the first 4 years of life, encompassing the juvenile period for females [median age at menarche in Amboseli = 4.5 years; 54]; (ii) had known mothers and fathers assigned using demographic and genetic data (see below); and (iii) had complete information on their experience of six sources of early-life adversity that together predict adult mortality [7, 51]: maternal loss, low maternal dominance rank, maternal social isolation, early-life drought, a close-in-age younger sibling, or large group size (see below). The females in our study were born into 13 different social groups, which are the fission or fusion products of two original study groups, first studied in 1971 and 1981.

### (b) Assigning maternities and paternities

The 216 female subjects were born to 117 mothers and sired by 102 fathers. Maternities were known from near-daily demographic records. Paternity assignments were based on microsatellite genotypes from at least six microsatellite loci and demographic records used to identify an initial pool of candidate fathers. These methods are described in detail in previous studies [30, 55–57], but briefly, microsatellite genotypes for juvenile females, mothers, and potential fathers were analyzed in the likelihood-based paternity assignment program CERVUS [58, 59]. We first included all potential fathers residing in the mother’s group at the time of conception (potential fathers are any male in the adult male hierarchy), and then expanded the set of potential fathers to include all adult, ranked males in the population. These two sets of analyses identified the same father in all but one case. In this case, we assigned paternity the sire who was seen consorting with the mother (the other potential sire lived in a different group and was never seen consorting the mother). Levels of confidence for all CERVUS analyses were set at 95%, and paternity assignments were robust across three rates of error, 1%, 5%, and 10% [30, 55–57].

### (c) Measuring grooming and co-residency between juvenile females and adult males

We defined co-residency as the cumulative number of days each juvenile female lived in the same group with her genetically confirmed father during the first 4 years of her life.

Following [6, 51, 60, 61], we measured annual pairwise grooming relationships for each year of the female’s life, using the “dyadic sociality index” (DSI). DSI provides a numeric score for each juvenile female’s dyadic relationship strength with individual adult males, scaled to be directly comparable to all other juvenile female-male pairs in the population in a given year of the female’s life, birthday to birthday. We calculated three types of DSI scores for juvenile females, which differed in the males included in the calculations: DSI_all_ measured dyadic relationships between juvenile females and all adult males who lived in her group for ≥30 days in the year in question, including her father (i.e., “co-resident” males); DSI_paternal_ measured dyadic relationships between juvenile females and their fathers; and DSI_non-paternal_ measured dyadic relationships between juvenile females and all co-resident males, excluding her father.

Because the ABRP only systematically collects focal animal samples on adult females and juveniles, and these data are sparse, DSI scores rely on grooming interactions collected via “representative interaction sampling” [6, 51, 60, 61]. During this sampling, observers record all grooming interactions between any interactants in their line of sight while simultaneously conducting random-order, 10-minute focal animal samples. These data are collected while observers continuously move throughout the group during focal sampling [6, 61], and prior work in our population finds that the resulting relative grooming frequencies are correlated with hourly rates of grooming from focal animal sampling [60].

From these data, we calculated each dyad’s log-transformed daily rate of grooming in a given year. Because dyads living in small groups experienced more intense representative interaction sampling than those in large groups, we accounted for differences in “observer effort” by regressing each dyad’s log-transformed daily rate of grooming against a measure of observer effort in that year: the number of focal animal samples per adult female per observation day for that group during the year in question. The resulting residuals were z-scored within years to estimate the DSI. Negative DSI scores were dyads who groomed less than was typical in that year; positive DSI scores were dyads who groomed more than was typical in that year. Note that the DSI_paternal_ score would be the same for a juvenile female who had no observed grooming with her father, whether they lived in the same group or not. We distinguish between these conditions by including both DSI_paternal_ and co-residency in our models of adult female survival (see below).

Our third objective required us to test why some fathers are more likely to groom their daughters than other males. For these analyses we compiled data on the presence or absence of male grooming directed to their daughters, in a given juvenile female year of life (contingent on ≥30 days of co-residency).

### (d) Measuring early-life adversity

To test if fathers moderate early-life effects on female mortality, we measured early-life adversity using a cumulative adversity index developed in prior studies [6, 7, 45, 51, 62]. This index sums the presence of six sources of early-life adversity: (1) drought in the first year of life (<200 mm of annual rainfall); (2) maternal death in the first four years of life; (3) being born into a large group as an index of realized resource competition (group size in the top quartile; ≥36 adults); (4) the birth of a close-in-age younger sibling that may divert maternal resources (interbirth interval in the shortest quartile, <1.5 years after the focal female’s own birth); (5) being born to a mother whose ordinal social dominance rank is in the bottom quartile for the population; and (6) being born to a mother who is in the top quartile for social isolation over the first 2 years of the juvenile’s life, measured based on an overall index of her involvement in grooming [7]. For each of the 216 juvenile females, we summed the number of these conditions that applied, resulting in a final index that could range from zero to six. No subject experienced more than four sources of adversity (20.3% of the 216 females experienced 0 sources of adversity; 40.2% experienced 1 source; 24.1% experienced 2 sources; 11.6% experienced 3 sources, and 3.7% experienced 4 sources).

### (e) Measuring predictors of father-daughter grooming and co-residency

For our third objective, we tested why some fathers are more likely to groom their juvenile daughters than other fathers. For a sub-set of variables, we also tested whether they explained the duration of father-daughter co-residency. Our sample sizes for these analyses were smaller than the 216 females in the first two objectives because we lacked information on some variables (see below).

*Male ordinal dominance rank* determines male priority of access to mates in our population, and mating opportunities could impose a tradeoff on time spent grooming offspring [63]. In Amboseli, male ranks are calculated monthly based on decided dyadic agonistic encounters between adult males [64].

The *daily rate of fertile females* in the group could also influence a male’s mating opportunities and impose a tradeoff on grooming offspring. This variable was calculated as the average daily number of peri-ovulatory females in the group in a given juvenile female-year on the days the male was resident in the group [65]. Peri-ovulatory periods are inferred from continuous records of sexual skin swellings that increase in size during the follicular phase and decrease during the luteal phase [65].

The *proportion of the mother’s available consort time the male obtained* during the 5-day peri-ovulatory period when the focal female was conceived predicts paternity [66] and possibly male paternity certainty. Hence, we summed all observed consort time that a mother had with any adult male within 5 days before the likely conception date of the focal female and calculated the proportion of this consort time that was monopolized by the male in question. Conception dates were calculated as described previously based on obvious signs of female reproductive state [65, 67]. For 31 of the 216 females, no males were observed consorting the focal female’s mother during her conceptive period. We excluded these 31 females because failing to observe a consort is much more likely to be caused by sparse behavioral sampling than the true absence of mate guarding.

*The number of potential fathers* present in the group at the juvenile female’s conception could influence male paternity certainty [56, 68]. This variable was calculated as the number of adult ranked males present in the group during the 5-day peri-ovulatory period when the female was conceived.

Following [34, 35], males who *sired the juvenile’s mother’s previous or subsequent offspring* might be more likely to groom their daughters if “primary associates” represent male mating effort. To test this possibility, we identified all cases in which the focal female’s father sired their mother’s previous or subsequent offspring. Including this variable led to further reductions in sample size because there were 42 cases where the paternity of either the female’s mother’s previous or subsequent infant was unknown.

The *number of offspring the male had in the group (i.e*., *co-resident offspring-years)* could influence his likelihood of remaining in his daughter’s group. This variable was calculated, for each father, as the number of his juvenile offspring that were alive in the group in a given juvenile female-year, scaled for days of co-residency. This variable may underestimate the male’s true count of living juvenile offspring because paternity is often missed for the youngest offspring (we generally obtain the first fecal sample between 6 and 18 months of age).

The female’s experience of *cumulative early-life adversity* (ELA), could also influence paternal investment. Cumulative ELA was calculated as the sum of the 6 conditions a female experienced prior to age 4 years (see above).

*Paternal age and the ages of daughters and their mothers* were known from near daily demographic records. All 216 daughters had ages accurate within a few days. Of the 102 fathers, 43 (42.2%) were born into the study population and their ages were accurate within a few days. For the remaining 59 fathers (57.8%), their ages were estimated to within a few years by comparing them to known-age males from the population [69]. For the mothers, 101 (86.3%) had ages accurate within a few days, 14 (12.0%) had ages accurate within 3 months, and 2 (1.7%) had ages accurate within 3 years.

For our model of why some fathers groom their daughters more than others, *we also modeled observer effort*, measured as the number of focal animal samples we collected per female-day (see above).

### (f) Statistical Analyses

Most analyses were performed in R 4.4.0 using the packages lme4 [70], lmerTest [71], lmtest [72], MuMIn [73], rptR [74], and survival [75]. We use an AICc-based information theoretic approach to test our hypotheses. See the GitHub repository cited in our data statement for a full list of packages.

#### Objective 1: Characterizing patterns of grooming and co-residency between juvenile females, their fathers, and other adult males

To measure grooming relationships between juvenile females and adult males, we calculated, in each female-year (i) the average number of adult males each female groomed with, and (ii) the percentage of grooming interactions she initiated with adult males (N=216 females; from 0 to 4 years of age). To test if female age predicted grooming initiation with males (both fathers and non-fathers), we ran a binomial LMM where the response variable measured whether grooming with adult males was initiated by females (1) or not (0), as a function of female age in each year of the juvenile period. Female identity was modeled as a random effect. To test if DSI_all_ was stronger between father-daughter pairs than other male-female pairs we compared Akaike information criteria (AICc) scores between two LMMs of juvenile females’ DSI_all_ scores: one that only included juvenile female age (0-4 years of age), and one that included juvenile female age and whether the male was the female’s father. Female identity was a random effect.

#### Objective 2: Testing if paternal co-residency and social bonds predict adult female survival

To test if females who have longer juvenile co-residency or stronger paternal grooming relationships exhibit higher adult survival than females who had short paternal co-residency or weak paternal grooming relationships, we ran a series of Cox proportional hazards models where the response variable was each female’s age at death or censorship, contingent on survival to her 4^th^ year of life (N=216 females; 124 censored values). Models were fit using coxph in the survival package [75]. We tested which variables best explained variation in adult female mortality risk based on AICc, including each female’s: (i) average DSI_paternal_ scores across the first four years of her life (because grooming patterns change with juvenile female age, these values were z-scored across females, within a year of life); (ii and iii) average DSI_non-paternal_ or DSI_all_ (also z-scored and averaged across the first four years of life); (iv) cumulative years of co-residency with her father in the first four years of life; and (v) cumulative early-life adversity score.

To test if the effects of early-life adversity on adult female survival are moderated by paternal grooming or co-residency, we added an interaction effect between females’ early-life adversity scores and (vi) their average annual DSI_paternal_ scores, and (vii) their cumulative years of paternal co-residency.

We also tested if juvenile females with strong paternal grooming bonds are socially well-connected with females and males in adulthood. If so, early-life relationships between female baboons and their fathers might be important for later life survival because they influence female social connectivity in adulthood, which predicts adult female survival in this population [51, 60, 61]. Social connectivity was calculated as a social connectedness index (SCI), which reflects the total amount of grooming the female, as an adult, gave and received with other adult females (SCI_F_) and adult males (SCI_M_) in her group [51, 60, 61]. For these analyses, we first ran an LMM testing if females with stronger mean DSI_paternal_ scores in the first 4 years of life had stronger SCI_F_ and SCI_M_ scores in adulthood, controlling for her age and dominance rank and modeling female identity as a random effect. We then ran a series of survival models to test if the association between DSI_paternal_ and adult female survival is attenuated by adding SCI_F_ and SCI_M_ to the model. The sample size for these models was 194 females because adult social connectedness information was missing for 22 females.

#### Objective 3: Testing the predictors of father-daughter grooming and co-residency

To test why some fathers are more likely to groom their daughters than others, we ran a binomial LMM where the response variable was whether a given father was observed to groom (1) or did not groom (0) his daughter in each of the first four years of her life, contingent on ≥30 days of co-residency. We used AICc to evaluate the predictive power of several fixed effects. These predictors included three indicators of a male’s mating opportunities in that year: (i) the father’s average dominance rank in that year, (ii) the average number of fertile (i.e., peri-ovulatory) females in the group each day in that year, (iii) and the number of other adult males (i.e., potential fathers) in the group at the daughter’s conception. As an indicator of paternity certainty, we used (iv) the proportion of observed consort time between the male and the focal female’s mother during the 5-day period when the female was conceived. As an indicator of whether male grooming of his daughter functioned as a form of mating effort with her mother, we included whether the father sired either the mother’s (v) prior or (vi) subsequent offspring. Because a male might invest less in any given daughter when he has many offspring, we controlled for (vii) the number of juvenile paternal offspring the father had in the group in that year (“co-resident offspring years”). We also controlled for (viii) the daughter’s cumulative ELA score, (ix-xi) the ages of the juvenile, the father, and the mother at the start of the year, and (xii) observer effort. Paternal identity was included a random effect (N=379 father-years involving 70 fathers and 130 juvenile females). We also performed a parallel analysis for all co-resident males (i.e., not just father-daughter pairs), which included a binary fixed effect for if the adult male was the father (N=6,324 father-years involving 288 males and 184 juvenile females with ≥30 days of co-residency).

To test why some fathers have longer co-residencies with their daughters than others, we ran an LMM where the response variable was the number of days the father resided in the same group as his daughter in the first four years of her life. The fixed effects: (i) the father’s dominance rank in the month co-residency ended; (ii) the daily rate of fertile females in the group in the month co-residency ended; (iii) the proportion of consort time the male had with the female’s mother; (iv) the number of other adult males (i.e., potential fathers) in the group at the daughter’s conception; (v) whether the father sired the mother’s previous offspring (subsequent offspring was excluded because only males with long co-residencies could sire future offspring); (vi) the number of juvenile paternal offspring the father had in the group in the year the co-residency ended (co-resident offspring years); and (vii) the daughter’s cumulative ELA score; and (viii-ix) the father’s and mother’s ages in the month co-residency ended (juvenile age was excluded because it was colinear with the duration of co-residency). Paternal identity was included as a random effect (N=166 co-residencies involving 86 fathers and 166 juvenile females).

Before performing our analyses, we checked for multicollinearity using variance inflation factor (VIF) analysis adapted for lmer models [76]. No variables had VIF >2.5.

## 3. RESULTS

### (a) Objective 1: Patterns of co-residency and grooming between juvenile females and their fathers

The median co-residency between the 216 daughters and their fathers was 33 months (**Fig. 1A**; range=0-48 months). More than a third of these females (n=80; 37%) lived in the same group with their father for ≥3 of their juvenile years. For the remaining 63% of females (n=136), their fathers either left the group or died sometime between their conception and 3 years of age (**Fig. 1A**). Thirteen females (6%) never co-resided with their fathers because the male dispersed or died between the focal female’s conception and birth.

**Figure 1.**
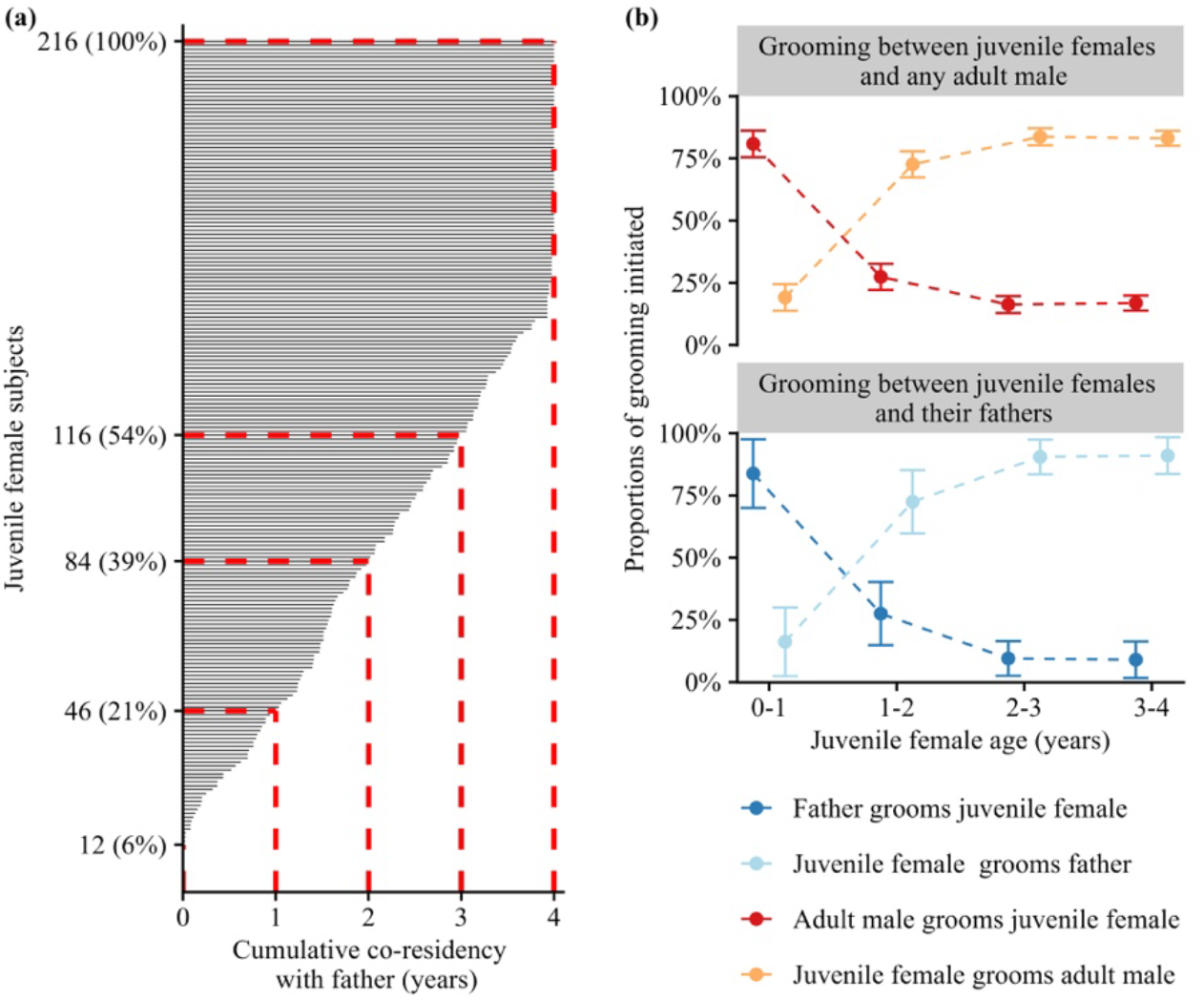
Co-residency and grooming directionality between juvenile females, their fathers, and other adult males. **(A)** Cumulative paternal co-residency (x-axis) for 216 juvenile females (y-axis). Each black bar represents the cumulative duration of time one female lived in the same group with her father. Red dashed lines demarcate the percentages of females who resided with their fathers for 1, 2, 3 or 4 years. **(B)** The average proportion of grooming interactions initiated by juvenile females (top: dark blue; bottom: red) or adult males (top: light blue; bottom: orange) as a function of female age. Top panel shows grooming initiation for fathers; bottom panel shows grooming initiation with all adult males.

Grooming between juvenile females, their fathers, and other adult males changed in frequency and directionality across females’ juvenile years. From birth to 4 years of age, females groomed with an increasing number of adult males (*β*=0.35; p<0.001) and were more likely to initiate grooming with adult males (**Fig. 1B**: binomial LMM: *β*=1.00; p<0.001). In the first year of life, however, 18.2% of grooming interactions with adult males were initiated by the females, and each female groomed with 1.15 adult males on average (range=1-3 males). By the 4th year of life, females had, on average, 1.54 male grooming partners (range=1-6 males), and 83% of these interactions were initiated by the female.

Consistent with prior evidence that males and their offspring have differentiated relationships [30, 32–36], daughters’ DSI_all_ values were stronger with their fathers than with other co-resident adult males (**Table 1;** ΔAICc for a model with and without “male is the father”=175.82). However, a visualization of DSI_all_ values between juvenile females and their fathers and between juvenile females and non-paternal males shows that some juvenile females also groom adult males who are not their fathers, and these bonds are sometimes strong (**Fig. S1**).

**Table 1.** LMM explaining dyadic bond strength between juvenile females and all co-resident adult males in a given year (N=10,833 DSI_all_ values between 216 females and 297 males, including 90 fathers).

| Term | $\beta$ (SE) | t | df | Interpretation |
| --- | --- | --- | --- | --- |
| Age of juvenile female | 0.152 (0.010) | 14.289 | 10133.1 | ↑juvenile age ↑bond strength |
| Male is the father | 0.696 (0.042) | 13.550 | 10105.4 | male is father ↑bond strength |

### (b) Objective 2: Early-life grooming and co-residency with fathers predicts adult female survival

We next tested whether daughters’ early-life relationships with their fathers predicted their adult survival. We found that juvenile females who had strong DSI_paternal_ scores with their fathers, or who had long co-residency with their fathers, or both, led longer adult lives than females with weaker paternal relationships (**Fig. 2**; **Table 2**). In support, the top three models predicting adult female survival included either the female’s average DSI_paternal_ score in the first four years of life, the duration of co-residency with her father, or both (**Fig. 2**; **Table 2 rows A-C**). DSI_paternal_ and co-residency were positively correlated with each other (**Fig. S2**; *β* =0.142, p<0.001), consistent with the idea that father-daughter pairs who co-reside for longer will also have stronger grooming relationships. Models that included either one or both variables were interchangeable in their ability to explain adult female mortality (**Table 2 rows A-C;** range in ΔAICc=0.31-1.28). This effect was specific to DSI_paternal_: strong relationships during the juvenile period with adult males in general (DSI_all_), or with non-fathers (DSI_non-paternal_), did not predict adult survival (**Table 2 rows A-C** versus **rows D, F, and G**).

**Table 2.**
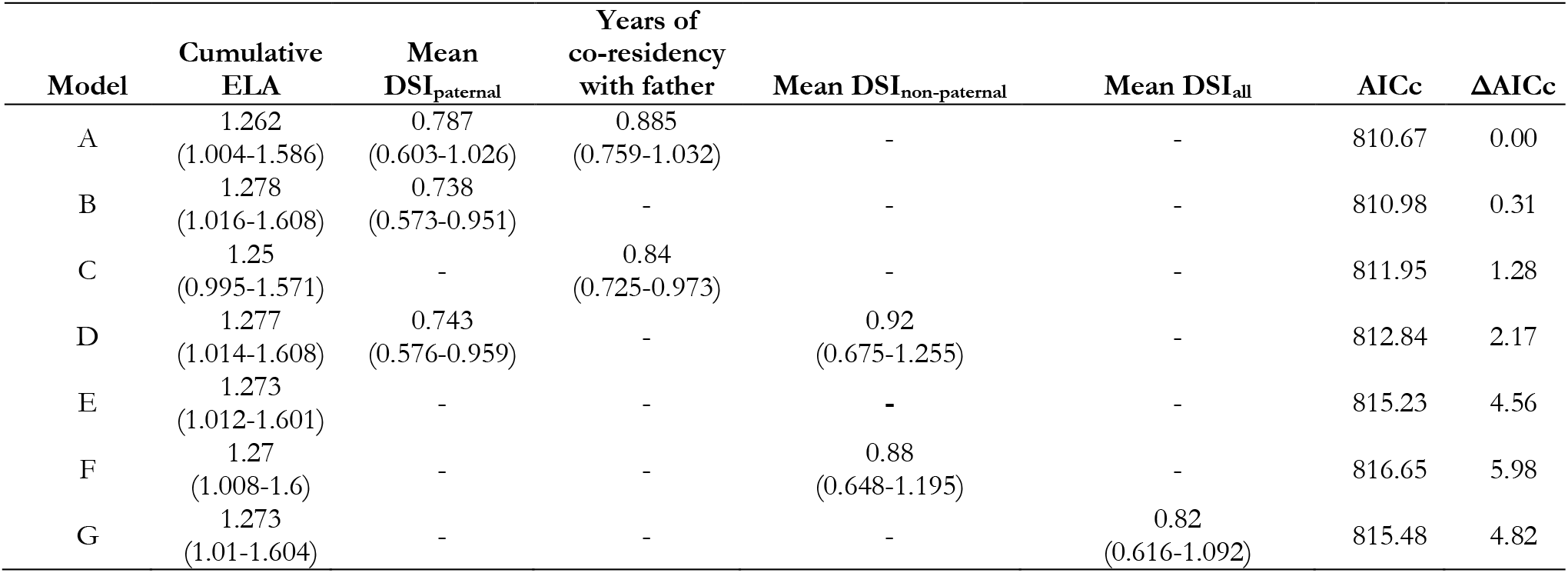
Results from seven alternative Cox proportional hazards models (n=216 females with 124 censored values) showing predictors of adult female survival. Each cell shows the variable’s hazard ratio (and 95% confidence interval). Models are ordered by AICc. No model violated the proportional hazards assumption.

| Model | Cumulative ELA | Mean DSI<sub>paternal</sub> | Years of co-residency with father | Mean DSI<sub>non-paternal</sub> | Mean DSI<sub>all</sub> | AICc | ΔAICc |
| --- | --- | --- | --- | --- | --- | --- | --- |
| A | 1.262<br>(1.004-1.586) | 0.787<br>(0.603-1.026) | 0.885<br>(0.759-1.032) | - | - | 810.67 | 0.00 |
| B | 1.278<br>(1.016-1.608) | 0.738<br>(0.573-0.951) | - | - | - | 810.98 | 0.31 |
| C | 1.25<br>(0.995-1.571) | - | 0.84<br>(0.725-0.973) | - | - | 811.95 | 1.28 |
| D | 1.277<br>(1.014-1.608) | 0.743<br>(0.576-0.959) | - | 0.92<br>(0.675-1.255) | - | 812.84 | 2.17 |
| E | 1.273<br>(1.012-1.601) | - | - | - | - | 815.23 | 4.56 |
| F | 1.27<br>(1.008-1.6) | - | - | 0.88<br>(0.648-1.195) | - | 816.65 | 5.98 |
| G | 1.273<br>(1.01-1.604) | - | - | - | 0.82<br>(0.616-1.092) | 815.48 | 4.82 |

**Figure 2.**
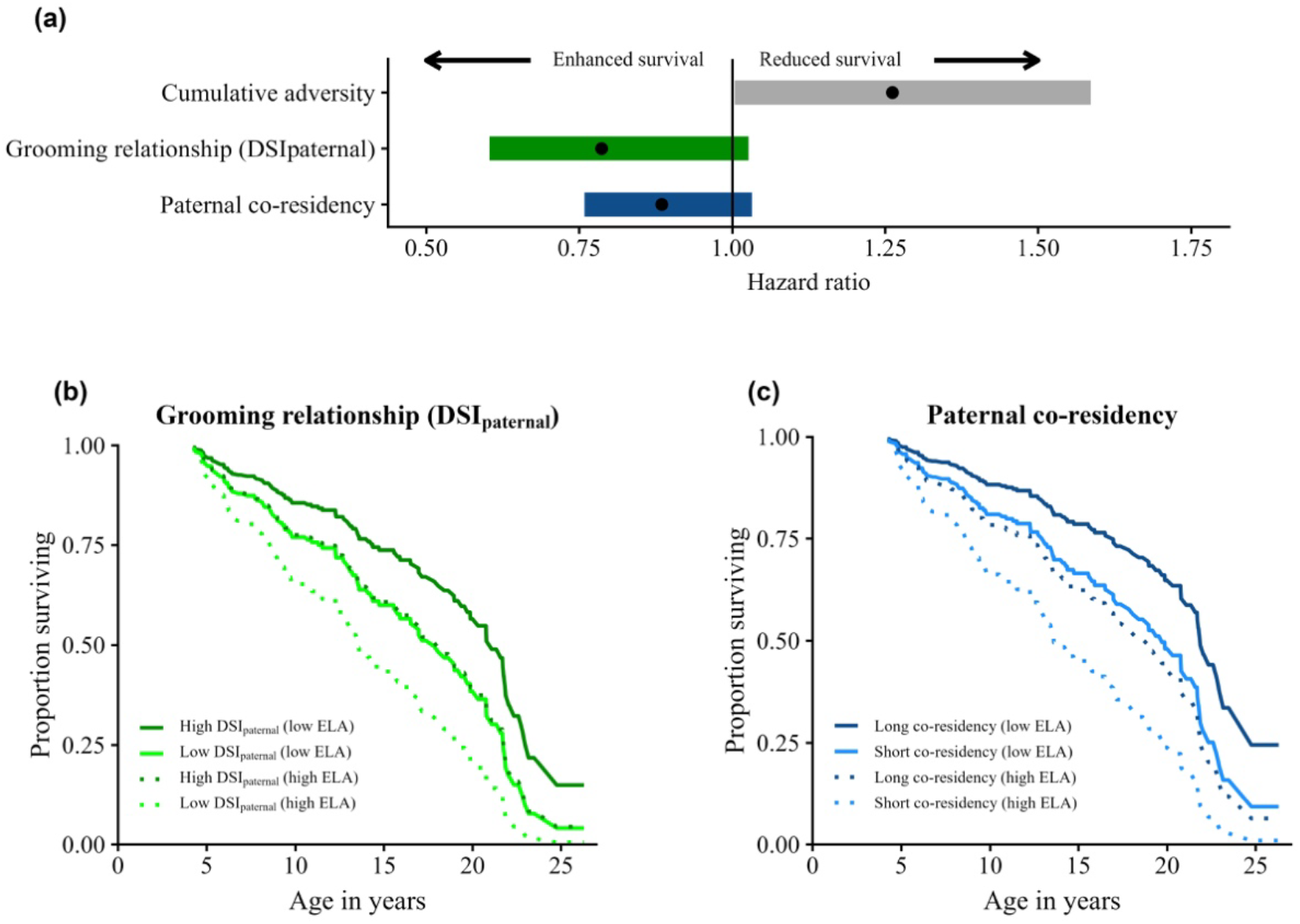
Juvenile females’ paternal grooming relationships and co-residency predict their adult survival. **(A)** Estimates from the time-to-event effects on the hazard of death in adult female baboons. Effects and 95% CI are from **row A in Table 2**, the model that had the lowest AICc and that includes mean DSI_paternal_ (green), paternal co-residency (blue) and early-life adversity (grey). **(B)** Predicted survival curves showing the effects of juvenile females’ mean DSI_paternal_ and early-life adversity (ELA) on adult female survival (predictions from model B in Table 2). Dark green lines are females in the top quartile of DSI_paternal_ scores; light green lines show females in the bottom quartile of DSI_paternal_. Solid lines show females who experienced one source of ELA; dashed lines show females who experienced 3 sources of ELA. **(C)** Predicted survival curves showing the effects of juvenile females’ duration of co-residency with their fathers and early-life adversity on adult female survival (predictions from model C in **Table 2**). Dark blue lines show females who lived with tsheir father for 1 year; light blue lines show females who lived with their fathers for all 4 juvenile years. Solid lines show females who experienced one source of ELA; dashed lines show females who experienced three sources of ELA.

Early-life adversity (ELA) also predicted adult female mortality (**Table 2 all models**, [7, 51]), but the models that included DSI_paternal_ and/or co-residency were a better fit to the data than a model that only included ELA (**Table 2 rows A-C vs row E**; range in ΔAICc=3.228-4.56). We therefore asked whether relationships with fathers predicted adult female survival more so for females who experienced harsh early-life circumstances. We found little evidence for an interaction effect between female ELA and either DSI_paternal_ or paternal co-residency (range in ΔAICc=1.66-3.57; **Table S1**). Hence, paternal relationships are not especially beneficial to high-adversity females. However, paternal relationships could still buffer females against some costs of ELA. Indeed, females who experienced three or more sources of adversity and were in the top quartile of paternal co-residency appear to survive as long as those who experienced just one source of adversity but had short paternal co-residency (**Fig. 2C**). For females who experienced one source of ELA, having a mean DSI_paternal_ score in the top quartile for the population predicted a median difference in survival of 1.8 years compared to females in the bottom quartile (**Fig. 2B**). Females who experienced one source of ELA and lived with their fathers for all 4 years were predicted to live 2.6 years longer than females who only lived with their father for 1 year (**Fig. 2C**). Females who experienced 3 or more sources of ELA were predicted to live 4.3-4.6 years longer if they co-resided with their father for 4 years versus 1 year or had grooming relationships with their fathers in the top versus bottom quartile (**Figs. 2B and 2C**).

We wondered whether some of the survival effects we observed could be explained by adult females’ social bonds with either sex in adulthood, both of which predict adult female survival [51, 60, 61, 77]. We found that females who had stronger average DSI_paternal_ scores across the first 4 years of life also had strong social bonds with adult males in adulthood (SCI_M_; **Table S2**). Females who had stronger average DSI_paternal_ scores also had stronger social bonds with adult females in adulthood, but in only one of the three best-supported models (SCI_F_; **Table S2**). In support of the idea that DSI_paternal_, SCI_F_, and SCI_M_ all contribute to adult mortality risk, four of the seven best-fitting models in **Table S3** included a metric of adult social connectedness (SCI_F_, SCI_M_, or both; **Table S3 models 3, 4, 6 and 7**), while six of the best seven models included DSI_paternal_, paternal co-residency, or both (**Table S3 models 1-6**).

### (c) Objective 3: Fathers are more likely to groom and live with their daughters when paternity is more certain and reproductive opportunities are limited

Father-daughter relationships should be stronger when males have greater paternity certainty and fewer reproductive opportunities. In support, males were more likely to groom their daughters in a given year if the male was low-ranking, if there were relatively few cycling females in the group that year (i.e., few reproductive opportunities), and if the male had a higher proportion of consort time with the female’s mother during the cycle the daughter was conceived (i.e., higher paternity certainty; **Table 3**; **Table S4**). Notably, we found that having more offspring in the group predicted more grooming, suggesting that males may increase their overall grooming time with juveniles if they have many offspring (**Table 3; Table S4**). Males who groomed their daughters were not more likely to sire the mother’s previous or next infant (**Table 3; Table S4**), consistent with the idea that grooming males are not investing in mating effort with the juvenile’s mother [33, 35, 36]. A similar subset of these variables also explained whether adult males had a grooming interaction with a juvenile female, regardless of whether the male was the father (**Table S5**), suggesting that a male’s rank and mating behavior at the time of an infant’s conception may influence his behavior towards that infant, regardless of whether he is the father.

**Table 3.** Model averaged estimates (*β*), standard errors (*SE*), and z-values for the seven best-supported GLMMs predicting the probability that a male groomed (1) or did not groom (0) his juvenile daughter in a given year of her life (N=379 father-years years involving 70 fathers and 130 juvenile females). Variables in bold appeared in all seven best-supported models (**Table S4**).

| Term* | $\beta$ ($SE$) | z-value | interpretation |
| --- | --- | --- | --- |
| **Father's average ordinal rank** | **0.185 (0.064)** | **2.868** | higher ordinal rank ↓grooming |
| **Daily rate of fertile females** | **-8.290 (2.425)** | **3.406** | ↑fertile females ↓grooming |
| **Proportion of consort time** | **2.334 (0.901)** | **2.581** | ↑consort time ↑grooming |
| Father sired the mother's next offspring | -0.949 (0.969) | 0.975 | Future offspring ↓grooming |
| Father sired the mother's previous offspring | -0.378 (1.001) | 0.376 | Previous offspring ↓grooming |
| **Father's co-resident offspring-years** | **0.235 (0.049)** | **4.822** | ↑co-resident offspring ↑grooming |
| Daughter's cumulative adversity score | -0.416 (0.391) | 1.061 | ↑adversity ↓grooming |
| **Juvenile age** | **0.778 (0.178)** | **4.362** | ↑juvenile age ↑grooming |
| Paternal age | 0.241 (0.201) | 1.195 | ↑male age ↑grooming |
| Maternal age | 0.035 (0.097) | 0.364 | ↑maternal age ↑grooming |
| **Observer effort** | **1.462 (0.247)** | **5.903** | ↑effort ↑grooming |

A slightly different set of variables predicted the duration of co-residency between fathers and their daughters (**Table 4; Table S6**). Fathers and daughters had longer co-residencies if the male was older, the mother was older, if there were more cycling females in the group, and if the male had more offspring in the group. Co-residencies also tended to be longer if the male had a prior offspring with the female’s mother.

**Table 4.** Model averaged estimates (*β*), standard errors (*SE*), and z-values for the five best-supported LMMs predicting the duration of father-daughter co-residency during the daughter’s 4-year juvenile period (N=166 co-residencies between 166 juvenile females and 86 fathers). Variables in bold appeared in all five best-supported models (**Table S6**).

| Term* | $\beta$ ($SE$) | z-value | interpretation |
| --- | --- | --- | --- |
| Daily rate of fertile females | 1.963 (1.030) | 1.892 | ↑fertile females ↑co-residency |
| Father sired the mother's previous offspring | -0.535 (0.239) | 2.226 | had offspring ↑co-residency |
| Father's co-resident offspring-years | 0.116 (0.036) | 3.228 | ↑offspring ↑co-residency |
| **Paternal age** | **0.160 (0.043)** | **3.672** | ↑male age ↑co-residency |
| Maternal age | 0.064 (0.020) | 3.086 | ↑maternal age ↑co-residency |

## 4. DISCUSSION

In many group-living mammals, males selectively interact with and provide low-cost forms of care to offspring [13, 22]. The selective pressures shaping these behaviors, and their importance to offspring health and survival, have received considerable attention in baboons [27, 30, 32–42]. Here we report that the strength of early-life paternal social relationships predicts meaningful differences in adult survival for female baboons in Amboseli, Kenya. These differences are on the order of 2-4 years for females in the top versus bottom quartile for paternal grooming or co-residency—effect sizes that are comparable to those for other major predictors of adult survival in Amboseli baboons, such as social isolation and early-life adversity [7, 61, 78]. This result joins prior evidence for early-life paternal effects in baboons, including the observations that paternal presence predicts earlier sexual maturity in daughters [32] and stronger social bonds between paternal half-siblings [44]. These early-life paternal effects may also be important in other mammal species that have subtle or indirect forms of paternal care [13, 17, 23–29]; hence, such effects may be more powerful and widespread than is currently appreciated.

### (a) What mechanisms explain early-life paternal social effects on female survival?

The two measures of father-offspring relationships we focused on—co-residency and grooming—are not themselves parental care, raising the question: how do these measures lead to early-life paternal effects? One possibility is that father-daughter grooming and co-residency are correlated with other male care behaviors, which in turn, directly benefit their daughter’s health and longevity. For instance, male baboons sometimes intervene on behalf of offspring in conflicts and buffer offspring and their mothers from negative interactions with other group members, including threats from infanticidal males [79–81]. These behaviors may reduce injury risk, improve offspring and maternal health, and ultimately affect offspring survival in adulthood [7, 62].

However, early-life relationships with fathers do not necessarily have direct, causal effects on offspring health and survival. Another possibility is that the strength of father-offspring relationships reflects daughters’ own phenotypic quality, which in turn, partly or completely explains the relationship between these traits and female survival. Under this scenario, strong, healthy juvenile females may build and maintain strong relationships with their fathers in early life, and also lead long, healthy adult lives. In support, father-daughter grooming relationships are largely maintained by daughters, not their fathers. Paternal effects may also be mediated by male health if paternal health affects offspring health, even in the absence of a direct relationship between fathers and offspring (through e.g., epigenetic marks or semen quality [18, 82]). Under this scenario, fathers who are in good condition produce healthy offspring, are less likely to disperse from groups where they fathered offspring, and are more socially engaged, including with their daughters. Such effects could create a correlation between father-offspring social relationships and offspring health and survival that is not directly causal.

Early-life paternal relationships could promote offspring social development [43], which in turn might affect survival via the established relationship between adult social connectedness and longevity [83]. In support, the juvenile females in our study were more socially connected in adulthood—especially to adult males—if they had stronger grooming relationships with their fathers. However, juvenile grooming with non-paternal males did not predict adult female survival— only grooming with fathers was protective. Further, paternal relationships predict adult survival, even controlling for adult social connectedness. Hence, the paternal effects we observed likely emerge from something more than adult social connectedness, and social connectedness to *all* males is likely not beneficial to females in the juvenile period, as it is in adulthood.

Much remains unknown about the mechanisms connecting father-daughter relationships to adult female longevity. One important next step is to test whether father-daughter grooming relationships are linked to more direct forms of care (e.g., carrying, agonistic support). We also do not yet know if fathers help their offspring survive the *juvenile* period, prior to adulthood. Further, data on daughters’ phenotypic quality and health in early life and adulthood are needed to test whether daughters with stronger paternal relationships are healthier and if so, which mechanisms mediate that effect (e.g., paternal care leading to offspring health, biological embedding of early experience, healthy females have more energy to pursue paternal bonds etc.).

### (b) Do male baboons experience selection to provide parental care?

If male care has direct effects on daughters’ lifespans and fitness, then our results help illuminate the evolution of paternal care. In Amboseli, adult female longevity is the largest contributor to female fitness for baboons, explaining ~80-90% of individual variation in adult female lifetime reproductive success [7, 45, 46]. Because female baboons typically produce one offspring every two years [84], the increased years of survival predicted by paternal co-residency or grooming could translate to 1-2 additional offspring over these females’ lifespans. As such, fathers and their juvenile daughters may experience selection to stay close and form lasting relationships.

However, male baboons are most likely to invest in social bonds with offspring when the reproductive tradeoffs are favorable [85–87]. For instance, the males in our population were more likely to groom their daughters when they had cues of paternity certainty—such as consorting with her mother—and few current mating opportunities. In practice, this meant that males were more likely to groom their daughters when they were low-ranking (because low-ranking males have low mating success) and/or when there were few fertile females in the group.

If males experience selection to remain in the same group with their daughters, the risk of inbreeding could increase. Indeed, the duration of father-daughter co-residency in our data indicates that baboons do experience this risk: 18% of females will co-reside with their father for some time period in adulthood [88]. However, even when fathers and daughters live together, mate guarding between father-daughter pairs in Amboseli is exceptionally rare [88]. Hence, father-daughter social relationships likely do not lead to costs of inbreeding for males or their daughters, and these relationships may in fact allow fathers and daughters to recognize each other and avoid mating.

We also found that males were more likely to have strong grooming relationships with their daughters and co-reside with their daughters if the male had more offspring in the group (“co-resident offspring years” in Tables 3 and 4). Hence, having more offspring does not appear to lead to a dilution in males’ tendencies to socialize with their daughters. While more research is needed to understand how being born into a large cohort of paternal siblings shapes baboons’ developmental environments, the lack of a dilution effect is consistent with the idea that, to the degree to which father-daughter grooming reflects parental care, most forms of parenting for male baboons are low effort, even if they carry substantial benefits to offspring.

These observations, together with the survival patterns we observed, suggest that baboon mothers may experience selection to increase males’ paternity certainty. Doing so would be advantageous to females if a prospective male sire meets other criteria for being a caring male (e.g., he sired a cohort of paternal siblings in the group). Selection for paternity certainty may also contribute to the evolution of sexual swellings in female baboons, which provide reliable signals of female ovulation and conception [35, 36, 65, 89, 90].

### (c) How do early-life paternal relationships interact with other early-life conditions?

We found little evidence for an interaction between females’ experiences of early-life adversity and the strength of their paternal relationships. As such, females who experienced harsh conditions in early life did not experience stronger survival benefits from paternal relationships than females who grew up in benign early-life environments. Hence, baboon fathers do not seem to actively fill deficits in their daughters’ early environments. However, despite the lack of an interaction effect, fathers may sometimes play important roles in buffering their daughters against the survival effects of early-life adversity. For instance, high-adversity females who have strong paternal grooming relationships or long paternal co-residency seem to survive just as long as low-adversity females with weak paternal relationships. These results parallel those for the effects of adult social connectedness and fecal glucocorticoid hormones in our population, both of which also have strong effects on adult survival, but appear to be mostly independent of early-life adversity [51, 78]. Hence, our data support the existence of multiple paths for mitigating early-life effects, including paternal relationships.

## ACKNOWLEDGEMENTS

We are grateful to Jeanne Altmann for stewarding the ABRP and in designing many of its data collection protocols. We acknowledge support from the National Science Foundation and the National Institutes of Health, currently through R01AG071684, R01AG075914, R01AG053330, R01AG053308, and R61AG078470. We thank the Max Planck Institute for Evolutionary Anthropology, Duke University, and the University of Notre Dame for financial and logistical support. In Kenya, our research was approved by the Wildlife Research Training Institute, Kenya Wildlife Service, the National Commission for Science, Technology, and Innovation, and the National Environment Management Authority. We also thank the University of Nairobi, the Kenya Institute of Primate Research, the National Museums of Kenya, the members of the Amboseli-Longido pastoralist communities, the Enduimet Wildlife Management Area, and Ker & Downey Safaris for their cooperation and assistance in the field. We thank T. Wango and V. Oudu for their assistance in Nairobi. The ABRP database is managed by J. Gordon and W. Wilbur, with contributions from N. Learn and K. Pinc. This research was approved by the IACUC at Duke University, University of Notre Dame, and the Ethics Council of the Max Planck Society. For further acknowledgments please visit http://amboselibaboons.nd.edu/acknowledgements/.

## DATA STATEMENT

Our data are publicly available on Zenodo at https://doi.org/10.5281/zenodo.14590285. Our code is available on GitHub at https://github.com/david-awam-jansen/BaboonPaternalRelationshipsSurvival

