## Supplementary materials for "Early-life paternal relationships predict adult female survival in wild baboons"

### **Supplementary Tables**

**Table S1.** Results from four Cox proportional hazards models (n=216 females) showing the effects of early-life adversity, paternal co-residency, and female bond strength with their fathers (DSI_paternal_), with interaction effects between early adversity included. Model A is identical to Model A in Table 2 in the main text (models B-G are also in Table 2). Each cell shows the hazard ratio for a given variable, with the 95% confidence interval in parentheses. Models are ordered by AICc. None of the models violated the Cox proportional hazards assumption (p-values ranged from 0.34 to 0.62).

| **Model** | **Cumulative early-life adversity** | **Mean DSI_paternal_** | **Years of**  **co-residency with father** | **Cumulative early-life adversity X Mean DSI_paternal_** | **Cumulative early-life adversity X years of**  **co-residency with father** | **AICc** | **ΔAICc** |
| --- | --- | --- | --- | --- | --- | --- | --- |
| A | 1.262 (1.004-1.586) | 0.787 (0.603-1.026) | 0.885 (0.759-1.032) |  |  | 810.67 | 0 |
| H | 1.28 (1.016-1.614) | 0.886 (0.591-1.328) | 0.883 (0.757-1.03) | 0.902 (0.683-1.192) |  | 812.33 | 1.66 |
| I | 1.212 (0.833-1.764) | 0.788 (0.604-1.028) | 0.863 (0.679-1.097) |  | 1.02 (0.883-1.178) | 812.78 | 2.12 |
| J | 1.174 (0.802-1.718) | 0.924 (0.601-1.421) | 0.833 (0.645-1.075) | 0.875 (0.649-1.179) | 1.045 (0.896-1.219) | 814.24 | 3.57 |

**Table S2.** Results from linear mixed models testing the relationships between a female’s social connectedness (SCI) to adult males (SCI_M_; upper table) and other adult females (SCI_F_; lower table) as a function of her average DSI_paternal_, her age, and her dominance rank. Female identity was modeled as a random effect. The upper and lower tables each show the four models with the lowest AICc; models within 2 AICc units of the best-supported model are shown in bold. Each cell shows the estimate given variable, with the 95% confidence interval in parentheses. The analysis includes 1,653 adult female-years for 194 females (social connectedness data are missing for 22 of the 216 females in our main data set).

| **Model** | **Mean DSI_paternal_ (in the juvenile period)** | **Female age at the start of a given year of adulthood** | **Average female dominance rank in a given year of adulthood** | **AICc** | **ΔAICc** |
| --- | --- | --- | --- | --- | --- |
| Predictors of adult female SCI_M_ | | | | | |
| **1** | **0.157**  **(0.079-0.235)** |  | **-0.019**  **(-0.029 - -0.009)** | **3,926.82** | **0** |
| **2** | 0.171  (0.090-0.252) |  |  | 3,929.47 | 2.65 |
| **3** |  |  | -0.020  (-0.031 - -0.010) | 3,935.12 | 8.31 |
| 4 | 0.157  (0.079-0.235) | 0.008  (-0.004-0.020) | -0.019  (-0.029 - -0.009) | 3,935.43 | 8.61 |
| Predictors of adult female SCI_F_ | | | | | |
| **1** |  |  | **-0.025**  **(-0.037 - -0.014)** | **3,821.73** | **0** |
| **2** |  | **-0.018**  **(-0.029 - -0.006)** | **-0.026**  **(-0.037 - -0.015)** | **3,822.84** | **1.11** |
| **3** | **0.107**  **(0.006-0.209)** |  | **-0.025**  **(-0.036 - -0.013)** | **3,823.54** | **1.82** |
| 4 | 0.110  (0.008-0.211) | -0.018  (-0.029 - -0.007) | -0.025  (-0.036 - -0.014) | 3,824.46 | 2.73 |

**Table S3.** Results from eight Cox proportional hazards models (n=194 females with 111 censored values) showing the effects of early-life adversity, paternal co-residency, Mean DSI_paternal_, SCI_F_, and SCI_M_ on adult survival. Models 1, 2, and 5 include the same variables as the three best-supported models in Table 2 in the main text (models A-C). Models 3, 4, 6, and 7 had ΔAICc <2 compared to model 1, but include SCI_F_ and SCI_M_ in adulthood. Each cell shows the hazard ratio for a given variable, with the 95% confidence interval in parentheses. Models are ordered by AICc.

| **Model** | **Cumulative early-life adversity** | **Mean DSI_paternal_** | **Years of**  **co-residency with father** | **SCI_F_*** | **SCI_M_*** | **AICc** | **ΔAICc** |
| --- | --- | --- | --- | --- | --- | --- | --- |
| 1 | 1.321 (1.033-1.688) | 0.745 (0.573-0.967) |  |  |  | 712.08 | 0 |
| 2 | 1.308 (1.023-1.671) | 0.788 (0.599-1.037) | 0.897 (0.762-1.055) |  |  | 712.49 | 0.41 |
| 3 | 1.373 (1.061-1.776) | 0.86 (0.641-1.153) | 0.888 (0.755-1.046) | 1.118 (0.76-1.642) | 0.665 (0.415-1.065) | 713.85 | 1.77 |
| 4 | 1.386 (1.069-1.797) | 0.805 (0.609-1.063) |  | 1.111 (0.762-1.62) | 0.677 (0.421-1.087) | 713.61 | 1.53 |
| 5 | 1.296 (1.015-1.656) |  | 0.85 (0.728-0.992) |  |  | 713.48 | 1.40 |
| 6 | 1.374 (1.063-1.777) |  | 0.862 (0.739-1.007) | 1.114 (0.756-1.641) | 0.607 (0.393-0.938) | 712.65 | 0.57 |
| 7 | 1.393 (1.074-1.806) |  |  | 1.102 (0.755-1.609) | 0.584 (0.378-0.903) | 713.93 | 1.85 |
| 8 | 1.314 (1.028-1.68) |  |  |  |  | 715.59 | 3.52 |

* Social connectedness metrics controlled for female age and dominance rank

**Table S4.** Model estimates for the seven best-supported linear mixed models predicting the probability that a male groomed (1) or did not groom (0) his juvenile daughter in a given year of her life (N=379 father-years years involving 70 fathers and 130 juvenile females with >30 days of co-residency). Variables are presented in the same order as Table 3 in the main text. Bolded variables appeared in all seven models. Variables not in bold appeared in one or two best-supported models. Models are ordered by AICc.

| Model* | **Father’s average ordinal rank** | **Daily rate of fertile females** | **Proportion of consort time** | Father sired the mother’s next offspring | Father sired the mother’s previous offspring | **Father’s co-resident offspring-years** | Daughter’s cumulative adversity score | **Juvenile age** | Paternal age | Maternal age | **Observer effort** | AICc | ΔAICc |
| --- | --- | --- | --- | --- | --- | --- | --- | --- | --- | --- | --- | --- | --- |
| 1 | **0.190** | **-8.357** | **2.287** |  |  | **0.234** |  | **0.846** |  |  | **1.467** | 1,463.65 | 0 |
| 2 | **0.174** | **-8.138** | **2.218** |  |  | **0.236** |  | **0.632** | 0.229 |  | **1.477** | 1,464.44 | 0.798 |
| 3 | **0.192** | **-8.321** | **2.372** |  |  | **0.231** | -0.390 | **0.834** |  |  | **1.429** | 1,464.71 | 1.060 |
| 4 | **0.183** | **-8.384** | **2.399** | -0.949 |  | **0.240** |  | **0.857** |  |  | **1.480** | 1,464.84 | 1.192 |
| 5 | **0.175** | **-8.079** | **2.189** |  |  | **0.234** | -0.450 | **0.592** | 0.258 |  | **1.435** | 1,465.21 | 1.566 |
| 6 | **0.191** | **-8.370** | **2.418** |  | -0.378 | **0.234** |  | **0.844** |  |  | **1.467** | 1,465.62 | 1.975 |
| 7 | **0.189** | **-8.372** | **2.339** |  |  | **0.234** |  | **0.813** |  | 0.035 | **1.473** | 1,465.64 | 1.989 |

* The number of potential fathers at conception was not included in any of the seven best-supported models based on AICc

**Table S5.** Model estimates for the four best-supported linear mixed models predicting the probability that a male groomed (1) or did not groom (0) a given juvenile female in a year of her life (N=6,324 father-years years involving 288 males (70 of the 288 were fathers) and 184 juvenile females with >30 days of co-residency). Variables are presented in the same order as Table 3 in the main text. Bolded variables appeared in all four models. Models are ordered by AICc.

| Model | **Male is the father** | **Father’s average ordinal rank** | **Daily rate of fertile females** | **Proportion of consort time** | Father sired the mother’s next offspring | **Father sired the mother’s previous offspring** | **Father’s co-resident offspring-years** | **Juvenile age** | **Paternal age** | Maternal age | **Observer effort** | AICc | ΔAICc |
| --- | --- | --- | --- | --- | --- | --- | --- | --- | --- | --- | --- | --- | --- |
| 1 | **0.648** | **0.053** | **-1.692** | **1.48** | -0.159 | **0.329** | **0.073** | **0.659** | **0.159** |  | **0.749** | 18008.31 | 0 |
| 2 | **0.638** | **0.053** | **-1.692** | **1.473** |  | **0.341** | **0.068** | **0.657** | **0.159** |  | **0.745** | 18009.39 | 1.08 |
| 3 | **0.648** | **0.053** | **-1.689** | **1.479** | -0.160 | **0.329** | **0.072** | **0.655** | **0.160** |  | **0.741** | 18009.57 | 1.26 |
| 4 | **0.648** | **0.053** | **-1.687** | **1.479** | -0.159 | **0.323** | **0.072** | **0.675** | **0.159** | -0.018 | **0.744** | 18009.58 | 1.27 |

* The female’s cumulative adversity score was not included in any of the seven best-supported models based on AICc

**Table S6.** Model estimates (*β)*, standard errors on the estimate (*SE*) and z-values for the five best-supported linear mixed models predicting the duration of father-daughter co-residency during the daughter’s 4-year juvenile period (N=166 co-residencies between 166 juvenile females and 86 fathers). Variables are presented in the same order as Table 4 in the main text. Variables in bold appeared in all five models.

| Model* | Daily rate of fertile females | Father had previous offspring with the mother | Father’s co-resident offspring-years | **Paternal age** | Maternal age | AICc | ΔAICc |
| --- | --- | --- | --- | --- | --- | --- | --- |
| 1 | 1.413 | -0.589 |  | **0.160** | 0.066 | 526.51 | 0 |
| 2 | 2.408 |  | 0.121 | **0.157** |  | 526.59 | 0.081 |
| 3 | 2.144 |  | 0.121 | **0.146** | 0.058 | 526.75 | 0.241 |
| 4 | 1.910 | -0.414 | 0.104 | **0.158** | 0.066 | 527.10 | 0.586 |
| 5 |  | -0.619 |  | **0.162** | 0.068 | 528.19 | 1.677 |

* The following variables were not included in any of the five best-supported models based on AICc: the daughter’s cumulative adversity score, the father’s ordinal rank, the proportion of consort time the father received, and the number of potential fathers at conception.

### **Supplementary Figures**

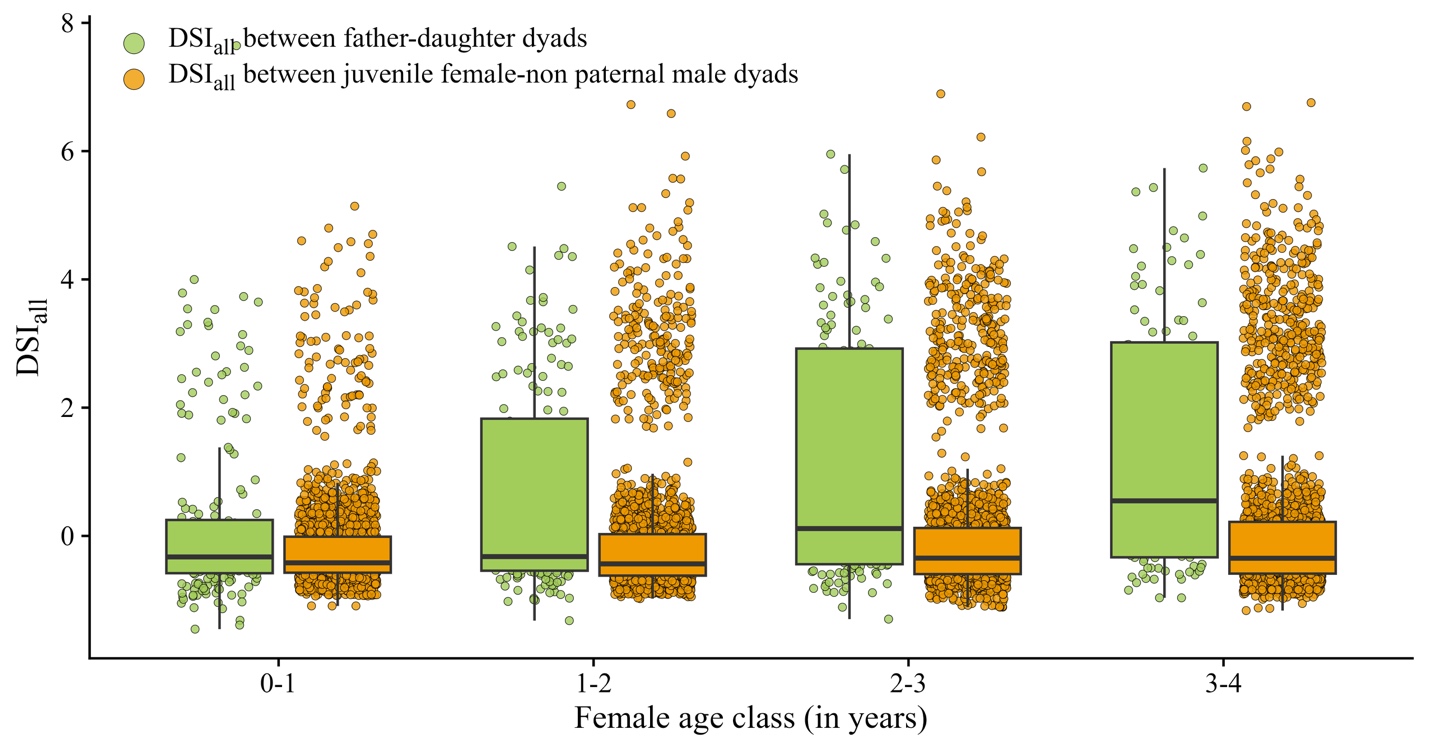

**Figure S1**. **DSI_all_ between juvenile females and all co-resident adult males as a function of female age in years.** Each point represents one DSI_all_ value for one year of co-residency for a father-daughter pair (green points; n=216 father-daughter pairs) or one DSI_all_ value for each year of co-residency between a juvenile female and one co-resident, non-paternal adult male (orange points; n=xx non-paternal pairs). Dyads who were not observed to groom in a given year of life were assigned a very low frequency of grooming so that they can be included in the dataset. This was necessary because grooming between juvenile females and adult males is uncommon, at least compared to grooming between mothers and daughters and between adult females.

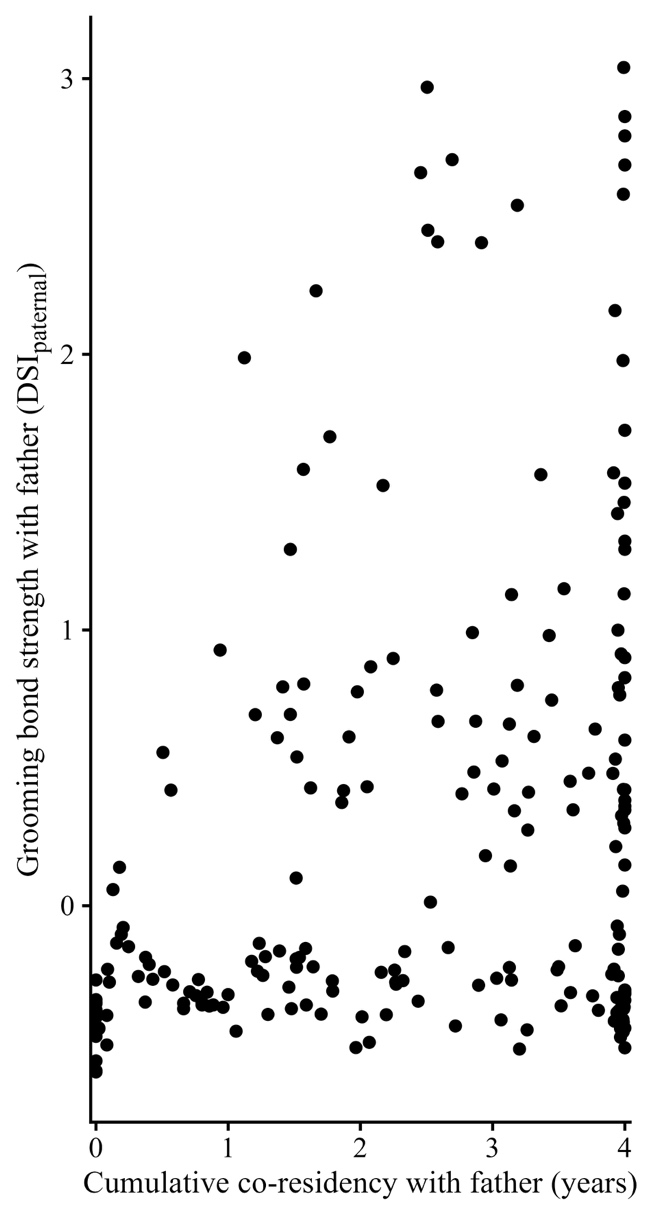

**Figure S2.** Father-daughter pairs with longer periods of co-residency had stronger DSI_paternal_ scores (*β* =0.142, p<0.001; Pearson’s correlation r=0.274). Each point represents one of the 216 father-daughter dyads in the data set.
